# A Live Attenuated Vaccine Candidate against Emerging Highly Pathogenic Cattle-Origin 2.3.4.4b H5N1 Viruses

**DOI:** 10.1101/2025.03.28.646033

**Authors:** Ahmed Mostafa, Chengjin Ye, Ramya S. Barre, Vinay Shivanna, Reagan Meredith, Roy N. Platt, Ruby A. Escobedo, Mahmoud Bayoumi, Esteban M. Castro, Nathaniel Jackson, Anastasija Cupic, Aitor Nogales, Timothy J. C. Anderson, Adolfo García-Sastre, Luis Martinez-Sobrido

**Author notes:** Correspondence should be addressed to: Luis Martinez-Sobrido, Ahmed Mostafa, PhD.

## Abstract

Influenza viruses present a significant public health risk, causing substantial illness and death in humans each year. Seasonal flu vaccines must be updated regularly, and their effectiveness often decreases due to mismatches with circulating strains. Furthermore, inactivated vaccines do not provide protection against shifted influenza viruses that have the potential to cause a pandemic. The highly pathogenic avian influenza H5N1 clade 2.3.4.4b is prevalent among wild birds worldwide and is causing a multi-state outbreak affecting poultry and dairy cows in the United States (US) since March 2024. In this study, we have generated a NS1 deficient mutant of a low pathogenic version of the cattle-origin human influenza A/Texas/37/2024 H5N1, namely LPhTXdNS1, and validated its safety, immunogenicity, and protection efficacy in a prime vaccination regimen against wild-type (WT) A/Texas/37/2024 H5N1. The attenuation of LPhTXdNS1 *in vitro* was confirmed by its reduced replication in cultured cells and inability to control IFNβ promoter activation. In C57BL/6J mice, LPhTXdNS1 has reduced viral replication and pathogenicity compared to WT A/Texas/37/2024 H5N1. Notably, LPhTXdNS1 vaccinated mice exhibited high immunogenicity that reach its peak at weeks 3 and 4 post-immunization, leading to robust protection against subsequent lethal challenge with WT A/Texas/37/2024 H5N1. Altogether, we demonstrate that a single dose vaccination with LPhTXdNS1 is safe and able to induce protective immune responses against H5N1. Both safety profile and protection immunity suggest that LPhTXdNS1 holds promise as a potential solution to address the urgent need for an effective vaccine in the event of a pandemic for the treatment of infected animals and humans.

**Graphical abstract:** 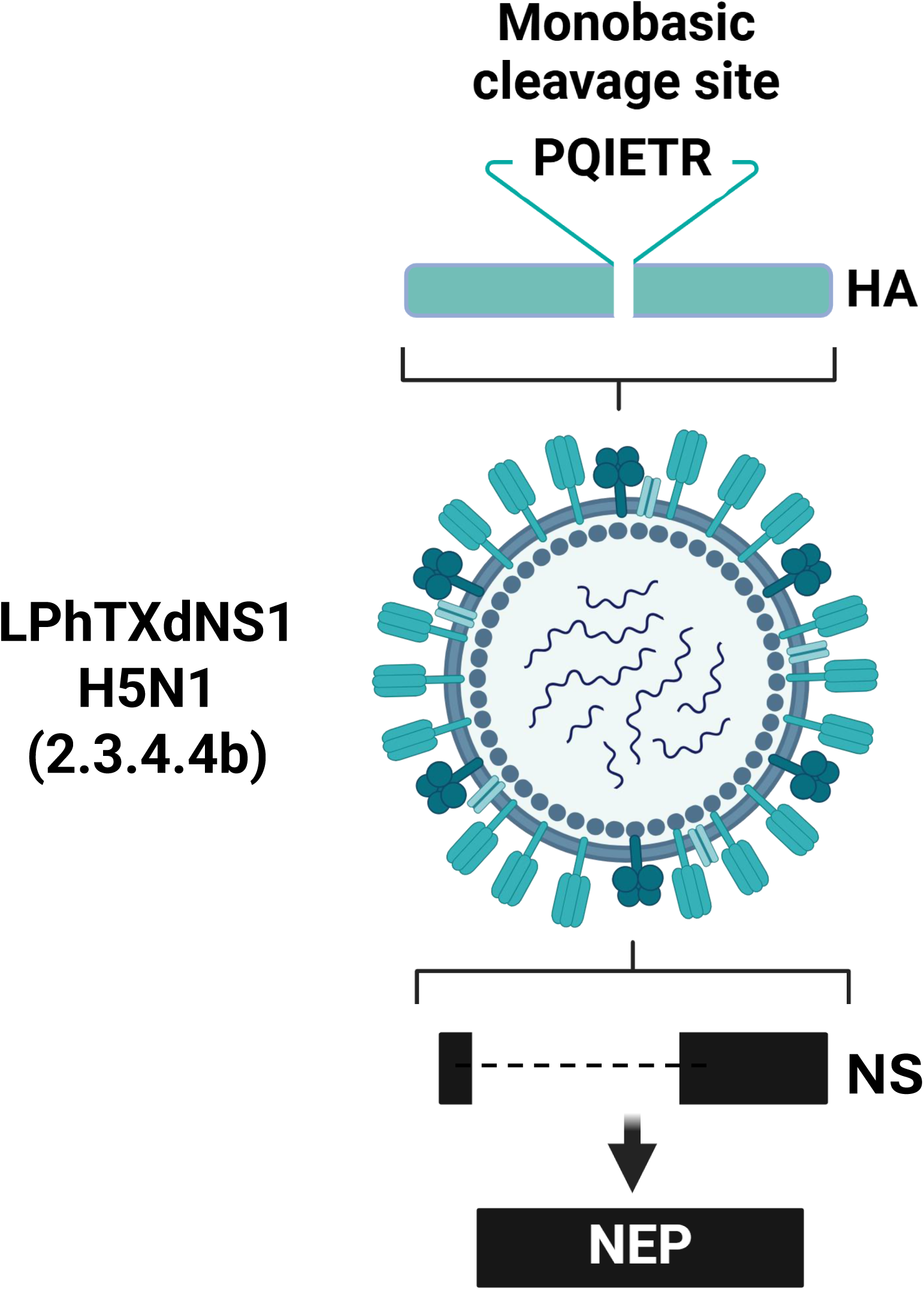

## Introduction

H5N1 avian influenza virus clade 2.3.4.4b emerged in 2020 and became dominant in Asia, Europe, and Africa by 2021. It spread to North America by the end of 2021, causing poultry outbreaks, and reached Central and South America in 2022, affecting farmed animals, mammals, and wildlife. In March 2024, a new genotype (B3.13) of H5N1 clade 2.3.4.4b emerged in the United States (US) to infect birds and dairy cattle, marking the first outbreak of H5N1 in ruminants (1). Shortly after, a human case was reported in a dairy farm worker in Texas following exposure to infected cattle, representing the first cattle-to-human transmission of H5N1 (1). Successively, H5N1 clade 2.3.4.4b genotype B3.13 resulted in massive outbreaks in cattle farms in the majority of the US, with remarkable increase in cattle-to-human transmission (1, 2). Amidst a yearlong H5N1 clade 2.3.4.4b genotype B3.13 outbreak in cattle farms, a second H5N1 clade 2.3.4.4b genotype D1.1 was also identified in early 2025 in cattle after transmission from birds (3).

Seasonal influenza vaccines reduce the burden of influenza disease, but their overall effectiveness varies each year (approximately 30-67%) (4–7), depending on how well the vaccine strain matches the circulating influenza viruses. Moreover, waning effectiveness has also been observed just a few months following vaccination, resulting in remarkable reductions in protection likely due to bias following the emergence of antigenic drift variants of influenza viruses that are not well-matched to the vaccine strains (7). In addition, the inactivated influenza vaccines induce a suboptimal immune response, especially when vaccine strains do not match circulating viruses, and fail to stimulate mucosal immunity, which could offer better protection against respiratory infections (8). Recent studies have shown that seasonal vaccination can boost antibody titers to protective levels in a moderate percentage of individuals against avian influenza viruses, likely due to cross-reactive immune responses (9, 10). These findings highlight the importance of vaccination and suggest that seasonal influenza vaccines could serve as an initial defense against a potential pandemic caused by avian influenza viruses, followed using more specific pandemic vaccines. Also, they highlight the significance of replacing the current inactivated vaccines with more effective live-attenuated influenza vaccines (LAIVs). LAIVs, which are recommended for individuals between the ages of 2 and 49, can trigger a range of immune responses by simulating a natural infection through intranasal administration, resulting in more robust and long-lasting immunity and protective effectiveness.

The current attenuated (*att*), temperature-sensitive (*ts*), cold-adapted (*ca*) 6+2 influenza virus reassortant seasonal LAIV based on the genetic background of influenza A/Ann Arbor/6/60 H2N2 (AA), has shown varying levels of effectiveness (11). The variable effectiveness of LAIVs is sometimes attributed to different factors, including improper selection of the vaccine strain, the attenuation method (12), or limitations in accurately assessing effectiveness due to limited use (11). The reduced effectiveness of seasonal LAIV in recent years may be also due to mismatching between the internal genes of the current LAIV strain, which are derived from viruses isolated between 1957 and 1960, and the currently circulating influenza viruses (13). Nonetheless, reports indicate that LAIVs can be up to 85% effective in children with a neutralizing antibody response durability for up to six months after vaccination (14–16). However, in adults, ca LAIV is poorly immunogenic, due to hyperattenuation of LAIV in the presence of immunity acquired by previous exposures to circulating influenza viruses (12, 17, 18).

The non-structural protein 1 (NS1) of influenza A virus (IAV) is crucial for interacting with many host proteins, suppressing the host’s antiviral response, and promoting viral replication and pathogenicity (19–21). IAV lacking NS1 are significantly weakened, making them viable for use as LAIVs (22). NS1-deficient (dNS1) LAIVs can mimic natural infection, induce robust immune responses, and protect against wild-type (WT) virus challenge, making them a promising option for emerging zoonotic avian influenza viruses (8). Recombinant avian, equine, canine, and swine IAV with partial truncations or deficiency in NS1 have been generated and proposed as potential LAIVs in diverse animal models including mice, guinea pigs, ferrets, chickens, horses, and dogs (13, 23).

Here, we investigated the impact of NS1 deletion on the replication, pathogenicity, virus-induced cellular and humoral responses, and protection efficacy of a low pathogenic cattle-origin human influenza A/Texas/37/34 H5N1 (LPhTXdNS1). Our results demonstrate that mice vaccinated with a single internasal dose of LPhTXdNS1 remain free of clinical signs of infection, exhibit strong immunogenicity, and protects against a subsequent lethal challenge with WT A/Texas/37/2024 H5N1, supporting its potential as an effective LAIV for combating the recent cattle-origin H5N1 highly pathogenic avian influenza virus (HPAIV) in the event of a pandemic.

## Materials and Methods

### Biosafety

All *in vitro* and *in vivo* studies involving highly pathogenic and low pathogenic avian influenza virus (HPAIV and LPAIV, respectively) H5N1 were conducted at biosafety level (BSL) 3 and animal BSL3 (ABSL3) laboratories at Texas Biomedical Research Institute (Texas Biomed). These experiments were approved by both the Texas Biomed Institutional Biosafety (IBC) and the Animal Care and Use (IACUC) committees.

### Cells

Madin-Darby canine kidney (MDCK), human embryonic kidney (293T), African green monkey kidney (Vero), and MDCK cells expressing GFP and firefly luciferase (FFluc) under the control of the interferon (IFN) β promoter (MDCK pIFNβ-GFP/IFNβ-FFluc) (24) were cultured in Dulbecco’s modified Eagle medium (DMEM) (Invitrogen, USA) complemented with 10% fetal bovine serum (FBS) and 1% PSG (penicillin, 100 U/mL; streptomycin, 100 µg/mL; L-glutamine, 2 mM). Cultured cells were kept at 37°C in a humidified 5% CO_2_ incubator.

### Plasmids and viruses

The recombinant low pathogenic human influenza virus A/Texas/37/2024 H5N1 (LPhTX) was generated as previously described (25) and propagated in MDCK cells using infection media (DMEM containing 1% P/S/G and 0.2% BSA) in the presence of 1 μg/mL of L-1-tosylamido-2-phenylethyl chloromethyl ketone (TPCK)-treated trypsin (Sigma–Aldrich, USA).

The pHW-hTX_NS plasmid (25), which expresses the NS1 and nuclear export protein (NEP) of A/Texas/37/2024 H5N1, was used as a template to remove the NS1-coding sequence. Briefly, 2 μl (5 ng/μl) of pHW-hTX_NS were combined with 5 μl of 10X Pfu reaction buffer, 1 μl of 10 mM dNTP mixture (0.2 mM of each), 1.5 μl of 50 mM MgCl2 (1.5 mM), 2 μl of forward primer (F: GTTAAGCTTTCAGGACATACTGATGAGGATGTCAAAAATGCAATTGGGGTCCTCAT) (40 pmoles), 2 μl of reverse primer (R: CTGAAAGCTTAACACAGTGTTGGAATCCATTATGTTTTTGTCACCCT) (40 pmoles), and 1 μl of PfuTurbo DNA Polymerase (2.5 U/μl) (Invitrogen, USA). The total volume was adjusted to 50 μl using nuclease-free water (Ambion, USA). The reaction was preceded by a pre-denaturation at 95°C for 2 min, followed by 35 PCR cycles (95°C for 30 sec for denaturation, 56°C for 30 sec for annealing, and 72°C for 3 min for extension), and a final extension step at 72°C for 10 min. The PCR product was purified using the Wizard® SV Gel and PCR Clean-Up System (Promega, USA). The resulting PCR product was then digested with HindIII (NEB, USA) to create the pHW-hTX-dNS1 plasmid.

To generate LPhTXdNS1, the seven ambisense pHW2000 plasmids containing the PB2, PB1, PA, monobasic HA, NP, NA, and M segments of LPhTX, and the pHW-hTX-dNS1 plasmid (1 μg each) were co-transfected into a 293T/MDCK cell co-cultures (3:1 ratio) using Lipofectamine™ 3000 Transfection Reagent (ThermoFisher Scientific, USA), according to the manufacturer’s protocol. After 24 h, the transfection medium was replaced with 1 mL of Opti-MEM media containing 1% PSG and 0.2% bovine serum albumin (BSA) and the culture was incubated in a 5% CO_2_ incubator. Twelve hours later, an additional 1 mL of Opti-MEM media containing 1% PSG, 0.2% BSA, and 2 μg/mL of TPCK-treated trypsin was added to each well. After 48-72 h, cell culture supernatants were harvested and centrifuged at 2,500 rpm for 5 min at 4°C. A portion of the collected supernatant was used to infect MDCK monolayers in T-75 cell culture flasks in infection media (DMEM with 1% PSG, 0.2% BSA, and 2 μg/mL of TPCK-treated trypsin). The rescued LPhTXdNS1 was then aliquoted and stored at −80°C.

Viral stock was confirmed by whole genome sequencing using the MinION platform (Oxford Nanopore Technologies) in comparison to the generated LPhTX strain. Briefly, viral RNA was extracted using the QiAamp Viral RNA Mini Kit (Qiagen, USA), and sample libraries were prepared with the Native Barcoding Kit 24 V14 (SQK-NBD114.24, Oxford Nanopore Technologies). Sequencing was performed on R10.4.1 Flow Cells (FLO-MIN114, Oxford Nanopore Technologies) according to the manufacturer’s instructions. Following nanopore sequencing, the initial read quality statistics were evaluated using n50 v1.7.0, followed by quality filtering with nanoq v0.10.0 (26). Reads were trimmed once average PHRED quality scores dropped below Q<10. In addition, we removed the first and last 15 base pairs of each read. After trimming, reads shorter than 500 base pairs were completely removed. Quality-trimmed reads were polished using the self-correcting algorithm LoRMA v0.4.2 (27) by comparing batches of 100 overlapping reads. Polished reads were aligned to the H5N1 LPhTX reference genome using minimap2 v2.28-r1209 (28) with the -x map-ont parameter to optimize mapping of error-prone, long Nanopore reads. Alignment statistics were calculated using SAMtools flagstat v1.21 (29), and genome-wide coverage was estimated using MosDepth v0.3.10 (30). We used LoFreq v2.1.5 (31) to evaluate indel quality and identify sequence variants. For variant calling, we capped the maximum allowable read depth at 10,000x (--max-depth) and disabled LoFreq’s default filters. Instead, we applied custom filtering criteria, retaining only variants supported by at least 25% of reads and located in regions with a minimum read depth of 100x.We also removed indels caused by homopolymer runs of three or more bases. For reproducibility, all analyses were completed on worker nodes containing 192 cores and 1Tb of memory at the Texas Biomedical Research Institutes high-performance computing center. All code necessary to complete the genetic analysis is available at GitHub (https://github.com/nealplatt/h5n1_highpathcow_2025-03-17). Sequence data is archived under BioProject accession code “PRJNA1242379”.

A recombinant A/Texas/37/2024 H5N1 containing a non-structural (NS) segment where the C-terminus of NS1 was fused to Nanoluciferase (NLuc) was engineered as previously described (32). The recombinant NS segment was synthesized *de novo* (Bio Basic, USA) with the appropriate restriction sites for subcloning into the ambisense plasmid pHW200 to generate the pHW-H5N1-NS_AgeI/NheI plasmid. The recombinant NS segment contained the NS1 open reading frame (ORF) without stop codons or splice acceptor sites, followed by AgeI and NheI restriction sites, the porcine teschovirus-1 (PTV-1) 2A autoproteolytic cleavage site (ATNFSLLKQAGDVEENPGP) and the entire ORF of the nuclear export protein, NEP (33). The Nluc ORF was cloned using the AgeI and NheI sites, into pHW-H5N1-NS_AgeI/NheI to generate the pHW-H5N1-NS_Nluc for virus rescue. Plasmid constructs were confirmed by DNA sequencing (Plasmidsaurus, USA). Recombinant A/Texas/37/2024(H5N1) virus expressing NS1_Nluc (HPhTX-Nluc) was rescued as previously described (25, 34–37). HPhTX-Nluc was then plaque purified and amplified on MDCK cells to generate the viral stock.

### Plaque Assay and Immunostaining

MDCK and Vero cell monolayers (6-well plate format, 10^6^ cells/well, triplicates) were infected with LPhTX and LPhTXdNS1 at multiplicity of infection (MOI) of 2. At 12 hpi, cells were washed with ice-cold PBS and treated with ice-cold NP40 lysis buffer (Thermo; completed with protease inhibitors cocktail). Cells were lysed on ice for 30 min, scrapped off and transferred to 1.5 mL Eppendorf tubes. Cells were centrifuged at 15,000 rpm for 15 min at 4°C. The cell lysate supernatants were then mixed with 2x working lithium dodecyl sulfate (LDS) sample buffer, supplemented with 20% β-mercaptoethanol and incubated at 98°C for 5 min before loading into SDS-PAGE gels. Following sample running, gels were transferred into nitrocellulose membranes. Subsequently, membranes were incubated in 5% non-fat dry milk powder in PBS with 0.05% Tween 20 blocking buffer (PBST) (Sigma, USA). Blocked membranes were probed overnight at 4°C with primary antibodies against PB1 (Clone F5-46, BEI Resources), PA (Clone 1F6, BEI Resources), NP (HT103) (38), NS1 (pAb, 1-73SW) (13), and β-actin (clone AC-15; A1978; Sigma, USA) as a house keeping protein. After overnight incubation, membranes were washed 3x for 5 min each with 5 mL of PBST. After washing, membranes were incubated with HRP-conjugated secondary antibodies for 2 h at RT. After 3x washes for 5 min with 5 mL of PBST, chemiluminescence-generated protein bands were visualized using a SuperSignal ECL substrate kit (ThermoFisher, USA) following manufacturer’s recommendations.

### Viral replication kinetics

Vero cell monolayers cultured in 6-well plates (10^6^ cells per well, triplicates) were infected with the specified viruses at a MOI of 0.001 and plates were kept at 37°C or 33°C in a humidified 5% CO_2_ incubator for 1 h to allow viral adsorption. After viral adsorption, the inoculum was removed and infected cell monolayers were washed 3x with PBS to eliminate any unadsorbed viral particles. Cell monolayers were then supplemented with 3 mL of infection medium containing 2 ug/mL of TPCK-treated trypsin and incubated at 37°C in a humidified 5% CO_2_ incubator. Aliquots of 200 µL were taken from the cell culture supernatants at 12, 24, 48, and 72 hpi and replaced with an equal volume of fresh infection medium. Viral titers in the collected samples were determined using plaque assays and immunostaining, as previously described (25, 39).

### Mice experiments

Female C57BL/6J mice (n=13/group) obtained from The Jackson Laboratory (JAX, USA) were anesthetized intraperitoneally (i.p.) by a cocktail of Ketamine (100 mg/mL) and Xylazine (20 mg/mL). Anesthetized mice were infected intranasally (i.n.) with the indicated viral doses in a total volume of 50 μl PBS. On 2-, 4-, and 6-days post-infection (DPI), 4 mice in each group were humanely euthanized to collect lung, nasal turbinate, and brain tissues. Half of the lung and brain organs were fixed in 10% neutral buffered formalin solution for histopathology and immunohistochemistry (IHC) analyses, and the other halves were homogenized in 1 mL of PBS using a Precellys tissue homogenizer (Bertin Instruments, USA) for viral titration. Tissue homogenates were centrifuged at 10,000 *xg* for 5 min and the supernatants were used to determine viral load by plaque assay and investigate the presence of secreted cytokines/chemokines. The remaining 5 mice/group were monitored for 14 days for body weight changes (morbidity), and survival rates (mortality). Mice that experienced a weight loss exceeding 25% of their original weight were humanely euthanized.

### Hemagglutination inhibition (HAI) assay

Sera were collected weekly from both mock-vaccinated and vaccinated mice (n = 5/group). The sera were treated with receptor-destroying enzyme (II) (RDE-II, Denka Seiken, USA) for 20 h at 37°C. The RDE-II enzyme was inactivated by heating at 56°C for 30 min. After treatment, 12.5 µl of the sera was added in triplicate to 96-well plates and mixed with 12.5 µl of diluted LPhTX, containing 4 hemagglutinating (HA) units. The plates were incubated for 30 min at RT. Then, 25 µl of 1% turkey red blood cells (RBCs) was added to each well, and after 45 min HAI was determined as the highest dilution of antiserum that completely inhibited hemagglutination (40).

### Histopathology and immunohistochemistry (IHC)

Mice lung and brain tissues were collected at necropsy, fixed in 10% neutral buffered formalin and processed in a Tissue Tek VIP tissue processor, where they were dehydrated through a series of graded alcohols, cleared with a 50:50 absolute alcohol/xylene mixture and two changes of xylene, and then infiltrated with paraffin wax using ParaPro™ XLT infiltration and embedding media (StatLab, USA). The paraffin blocks were sectioned at 4 microns thick using a Microm HM325 rotary microtome and mounted on microscope slides using a flotation water bath set between 46° and 48°C. The slides were then placed in a Varistain Gemini automated slide stainer for H&E staining, followed by drying in heating stations before the staining program began. The deparaffinization of tissue sections was done with three changes of xylene, two changes of absolute alcohol, and two changes of 95% alcohol, followed by rinsing in distilled water. Hematoxylin staining was conducted with Reserve Bluing Reagent (StatLab, USA), with excess stain removed using High-Def solution (StatLab, USA). The tissue sections were then stained with Eosin (StatLab, USA), dehydrated in three changes of alcohol, a 50:50 absolute alcohol/xylene mixture, and cleared in three changes of xylene before mounting them in coverslips. After staining, the tissue sections were examined by a board-certified veterinary pathologist in a blinded fashion. For IHC, tissue sections were cut at 4 microns, mounted onto positively charged slides, and allowed to air dry overnight. The slides were then processed using the Discovery Ultra IHC/ISH automatic stainer to detect avian influenza virus. Deparaffinization was performed using Discovery Wash (Roche, USA). Cell conditioning was done with Discovery CC1 (Roche, USA) at 95°C for 64 min. Endogenous peroxidase was blocked with Discovery Inhibitor (Roche, USA) for 8 min. Slides were then incubated with a rabbit polyclonal antibody against IAV NP (Invitrogen, USA) at a 1:1,500 dilution for 1 h at RT, followed by detection with an anti-rabbit HQ (Roche, USA) for 8 min, and anti-HQ HRP (Roche, USA) for 8 min at 36°C. Influenza NP was visualized using ChromoMAP DAB (Roche, USA). The slides were counterstained with Hematoxylin (Roche, USA) and blued using Bluing Reagent (Roche, USA). All HE stained and IHC stained slides were scanned using Zeiss axio scan. Z1 whole slide scanner at ×20 magnification. The digital slides were analyzed using an AI optimized tissue classifier in HALO v4.0 software (Indica Labs, USA) for absolute quantification of the percentage of lung affected and percentage of lung staining for viral antigen.

### Statistical analyses

All graphs, calculations, and statistical analyses were performed using GraphPad Prism software version 9.5.1 (GraphPad Software, LLC, USA). No assumptions of equal variance or sphericity were made for group comparisons. Normality of residuals was checked for all group comparisons and log10 transformations were applied when the assumption of normality was not met. Growth kinetics were analyzed using a two-way repeated measure ANOVA with Geisser-Greenhouse correction. Post-hoc comparisons were performed using the Šídák method. Differences in FFluc expression were evaluated using Welch’s one-way ANOVA, followed by multiple comparisons using Dunnett T3 method. Viral replication and titer data were analyzed using two-way repeated measure ANOVA with Geisser-Greenhouse correction. Post-hoc testing was conducted with Dunnett’s multiple comparisons test. Differences in survival curves were analyzed using a Log-rank test.

## Results

### Generation and characterization of LPhTXdNS1

IAV NS1 has been previously described to inhibit host gene expression, including type I interferon (IFN-I) responses and, therefore, contribute to viral virulence (20, 21). To establish an attenuated recombinant low pathogenic influenza A/Texas/37/2024 H5N1 with the potential to be implemented as a LAIV, we conducted virus attenuation by removing two key virulence viral factors, the NS1 protein and the multibasic cleavage site (PLREKRRKR/GLF) in the viral HA protein (**Fig. 1a**). Using our previously described reverse genetics for influenza A/Texas/37/2024 H5N1, we generated LPhTXdNS1, a virus lacking the NS1 protein and expressing a monobasic cleavage site (PQIETR/GLF) in the HA protein. We also generated a control A/Texas/37/2024 H5N1 virus, which has a monobasic cleavage site in its HA protein, but with an intact NS1 protein (LPhTX) (**Fig. 1a**).

**Figure 1.**
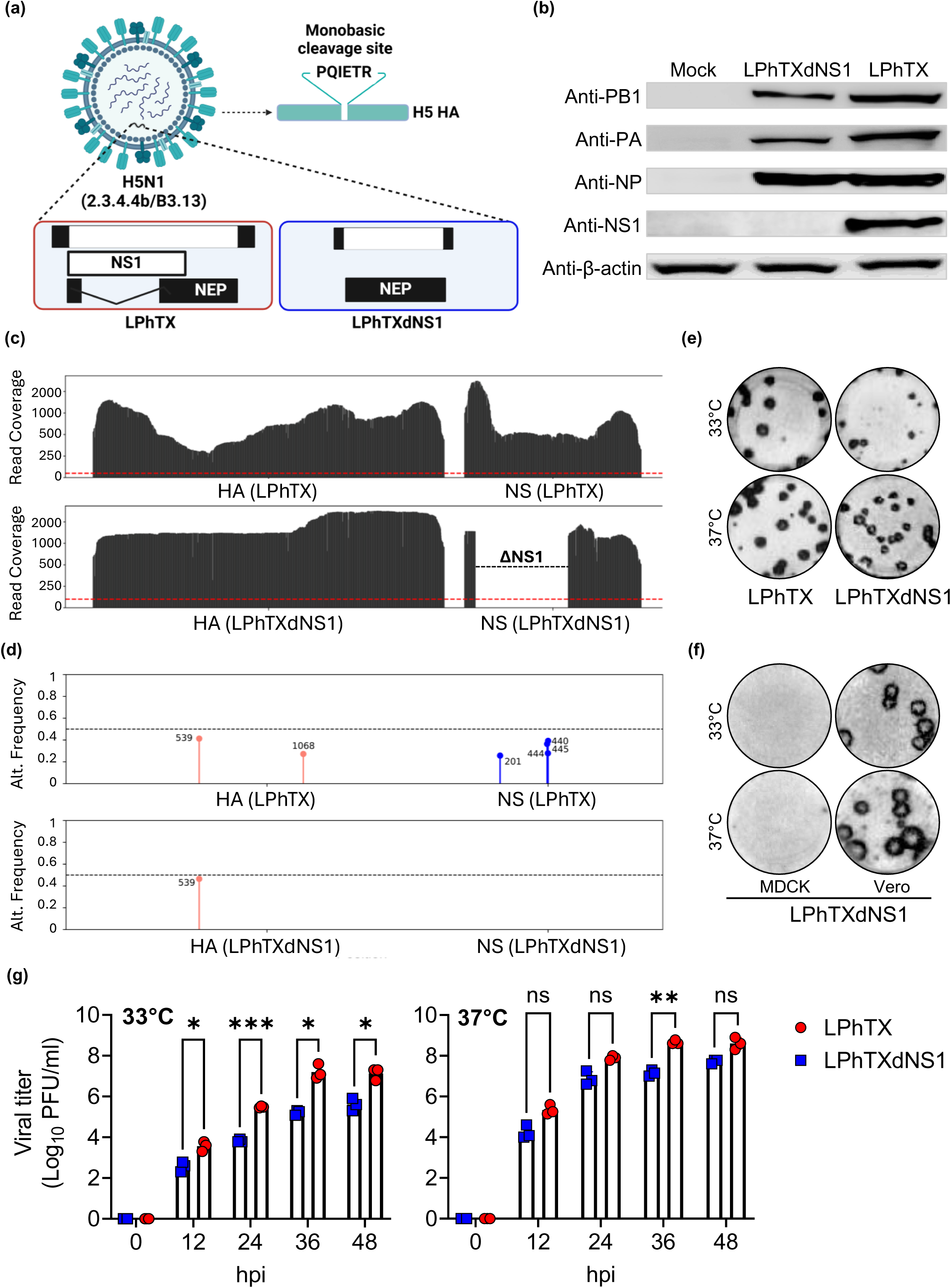
Generation and characterization of LPhTXdNS1. (**a**) Schematic representation of influenza low pathogenic human A/Texas/37/2024 H5N1 (LPhTX, left) and NS1 deficient (LPhTXdNS1, right) viruses. Mutation of the multibasic cleavage site of A/Texas/37/2024 H5N1 HA (PLREKRRKR/GLF) to a monobasic cleavage site (PQIETR) that made the backbone of LPhTX and LPhTXdNS1 is indicated on the top right. (**b**) Western blot showing the lack of NS1 protein expression in LPhTXdNS1-infected, compared to LPhTX-infected Vero cells. The PB1, PA, and NP viral proteins were included as viral infection controls. β-actin was included as a loading control. (**c**) Read depth across HA and NS segments – Coverage was calculated across the HA and NS segments for the LPhTX and LPhTXdNS1. The red dotted line represents 100x read depth. The NS gene in LPhTXdNS1 contains a complete NS1 deletion. (**d**) Non-reference allele frequency - The genome of the rescued viruses were compared to the LPhTX reference to identify variant sites. SNV allele frequency from LPhTX and LPhTXdNS1 are shown with circles. Variants on the HA and NS segments are shown in red and blue respectively. Variants <25% frequency and indels caused by homopolymer runs greater than 3 nt are not shown. Allele frequencies are provided in Table 2. The red line indicates 50% allele frequency. (**e**) Plaque phenotypes of LPhTX and LPhTXdNS1 at 33°C (top) and 37°C (bottom) in Vero cells. Both viral stocks were titrated using immunostaining assay 24 hpi. (**f**) Plaque phenotypes of LPhTXdNS1 at 33°C (top) and 37°C (bottom) in MDCK (left) and Vero (right) cells by immunostaining at 48 hpi. (**g**) Growth kinetic of LPhTX and LPhTXdNS1 in Vero cells. Cell monolayers were infected at MOI=0.001 and viral titers at 12, 24, 36 and 48 hpi were determined by plaque assay using an immunostaining assay. A two-way repeated measure ANOVA with Geisser-Greenhouse correction. Post-hoc multiple comparisons performed using Šídák method to compare groups within each time-point. The significant differences are indicated (* = *p* < 0.05, ** = *p* < 0.01, *** = *p* < 0.001).

After rescuing LPhTXdNS1, the lack of NS1 expression was confirmed by Western blot (**Fig. 1b**) and next generation sequencing (**Fig. 1c**). In LPhTXdNS1-infected Vero cell lysates, the polymerase subunits PB1 and PA, and the viral NP, were detected at similar levels using specific antibodies in both LPhTXdNS1- and LPhTX-infected cells (**Fig. 1b**). However, NS1 expression was undetectable with an anti-NS1 antibody in Vero cell lysates infected with LPhTXdNS1 (**Fig. 1b**), contrary to LPhTX-infected Vero cells where NS1 expression was readily detectable.

On the same hand, the long-read sequencing data generated from Nanopore runs for LPhTX and LPhTXdNS1 were summarized in **Table 1**. In total, we analyzed 131.1 megabases (Mb). Read depth ranged from 427x-889x (**Table 1**; **Fig. 1c**). Reduced coverage in the LPhTXdNS1 gene was due, in part, to the dNS1 deletion. We identified 21 variants in the HA and NS segments after applying filtering criteria that removed variants with allele frequencies below 25% (**Table 2**; **Fig. 1d**). Of these, 14 were indels associated with homopolymer regions greater than 3 nt in length which are known to pose challenges for accurate sequencing using Nanopore technology (41). The frequencies of the remaining 7 variants were below 50% in each sample (**Table 2**). Among these seven non-silent variants, six (HA: G539A, C1068T; NS: C201G, C440G, C444T, and G445A) were associated with LPhTX, while only one variant (G539A) was linked to LPhTXdNS1.

**Table 1.** Data description. Sample summary and genome coverage

| Sample | n Reads | n50 | Mb | Align. Rate | Mean Coverage | >100x |
| --- | --- | --- | --- | --- | --- | --- |
| LPhTX | 78,248 | 1,183 | 104.1 | 98.89% | 426.6 | 99.99% |
| LPhTXdNS1 | 30,442 | 1,146 | 26.7 | 98.43% | 888.5 | 50.68% |

**Table 2.** Variant frequencies. All variants at <25% allele frequency are shown on the HA and NS segments. Homopolymer indels >3nt have been removed as well.

| Virus | Seg. | Pos. | Ref. allele (aa) | Alt. allele (aa) | Alt. freq | Read depth |
| --- | --- | --- | --- | --- | --- | --- |
| LPhTX | HA | 539 | G (Aspartic acid) | A (Asparagine) | 41.3% | 603 |
| LPhTX | HA | 1068 | C (Alanine) | T (Valine) | 27.2% | 1,380 |
| LPhTX | NS | 201 | C (Arginine) | G (Glycine) | 25.7% | 966 |
| LPhTX | NS | 440 | C (Phenylalanine) | G (Leucine) | 36.4% | 918 |
| LPhTX | NS | 444 | C (Arginine) | T (Tryptophan) | 27.8% | 916 |
| LPhTX | NS | 445 | G (Arginine) | A (Glutamine) | 39.2% | 921 |
| LPhTXdNS1 | HA | 539 | G (Aspartic acid) | A (Asparagine) | 46.5% | 863 |

The plaque sizes of LPhTXdNS1 in Vero cells at 33°C and 37°C was slightly smaller than LPhTX, containing a functional NS1 protein (**Fig. 1e**). To confirm that LPhTXdNS1 is limited in its ability to replicate in immunodeficient but not in immunocompetent cells, we conducted plaque assays of LPhTXdNS1 in MDCK and Vero cells, respectively, at 33°C and 37°C for 48 hpi. No viral plaques were detected in MDCK cells, whereas LPhTXdNS1 was able to form plaques in Vero cells (**Fig. 1f**). As expected, the LPhTXdNS1 replicates in IFN-I deficient Vero cells, reaching viral peaks at ∼48 hpi to average titers of 1.7×10^5^ and 1.467×10^7^ PFU/ml at 33°C and 37°C, respectively (**Fig. 1g**) that is ∼30 and ∼12 times lower than LPhTX at same corresponding temperatures, respectively. Altogether, these results demonstrate that NS1 deletion in LPhTXdNS1 renders the virus attenuated *in vitro* when compared to the NS1-expressing LPhTX virus.

### LPhTXdNS1 lacks the ability to inhibit IFNβ promoter activation

We further confirmed the lack of NS1 expression by LPhTXdNS1 by infecting MDCK cells constitutively expressing GFP-CAT and FFluc under the control of an IFNβ promoter (24), designated hereafter as MDCK IFNβ GFP-CAT/FFluc (**Fig. 2a**). As expected, LPhTX was able to efficiently control IFNβ promoter activation as determined by the lack of GFP and reduced FFluc expression (**Figs. 2b-c**). However, cells infected with LPhTXdNS1 showed high levels of GFP and FFluc expression, compared to LPhTX, due to the lack of NS1 expression (**Fig. 2b-c**). The results further emphasize that the LPhTXdNS1 is ineffective in inhibiting IFNβ promoter activation in immunocompetent MDCK cell lines, causing the virus to be vulnerable to the antiviral activity of IFN that hinders its efficient replication in these immune-competent cell culture models.

**Figure 2.**
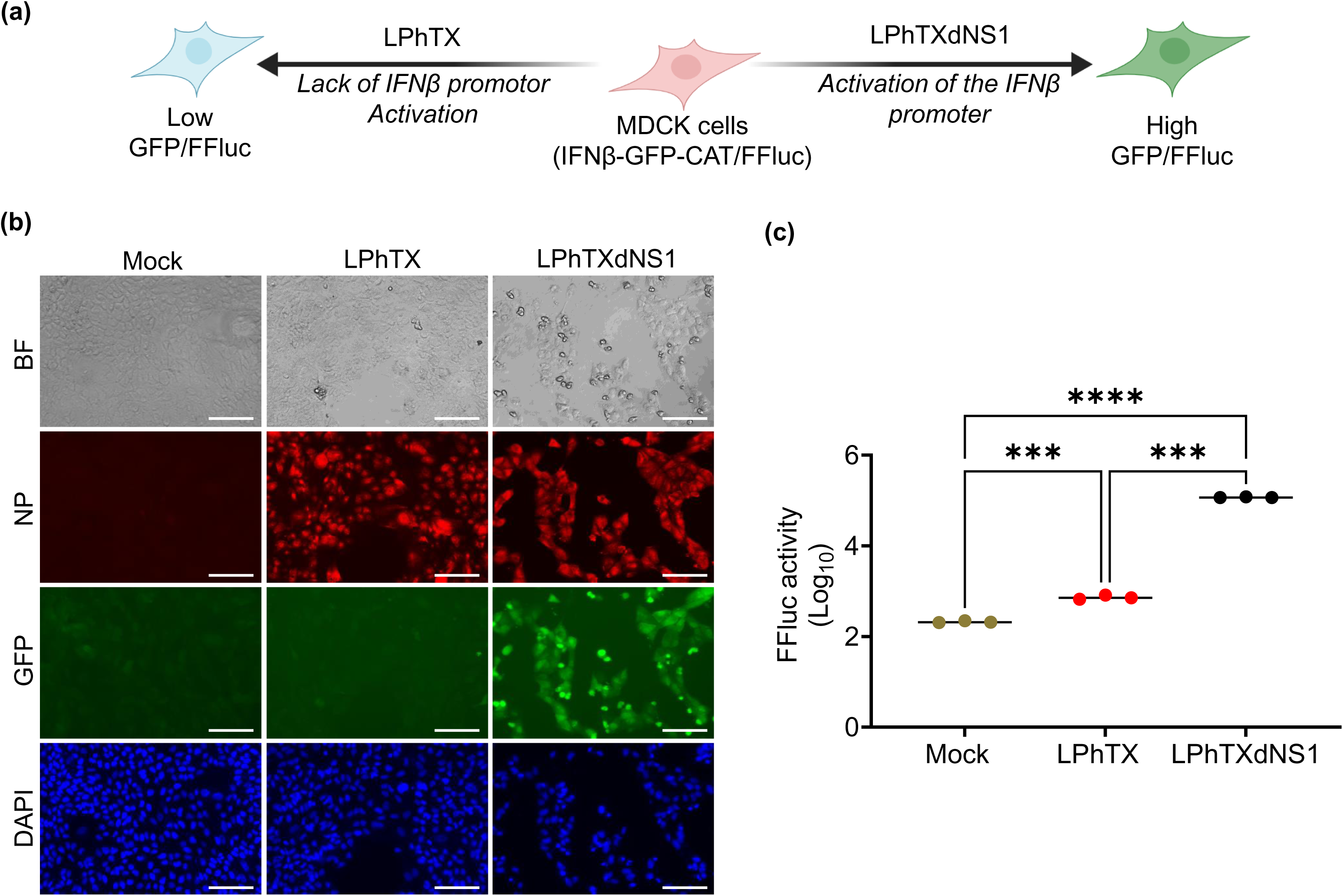
IFNβ promoter activation by LPhTXdNS1 infection. (**a**) Schematic representation of the IFNβ induction bioassay: MDCK cells constitutively expressing GFP-CAT and FFluc reporter genes under the control of the IFNβ promoter (MDCK IFNβ GFP-CAT/FFluc) were infected at MOI 2 with LPhTX, LPhTXdNS1, or mock-infected. At 12 hpi, IFNβ promoter activation was determined by GFP (**b**) and FFluc (**c**) expression. The viral NP in infected cells was detected with the HT103 MAb and DAPI was used for nuclear staining. The scale bar was set for 100 μm. BF: Bright-Field; GFP: Green Fluorescent Protein. All data points are shown with black lines representing the mean (*n* = 3). Groups compared using a Welch’s one-way ANOVA, followed by multiple comparisons using Dunnett T3 method (*** = *p* < 0.001; **** = *p* < 0.0001).

### Pathogenicity and immunogenicity of LPhTXdNS1

To determine the safety of LPhTXdNS1, female C57BL/6J mice were infected (i.n.) with 10^2^-10^4^ PFU of LPhTXdNS1. Uninfected mice and mice infected with 10^2^ PFU of LPhTX were included as controls. Mice were monitored daily for morbidity and survival up to 14 DPI (**Fig. 3**). Mice infected with 10^2^ PFU of LPhTX, where the multibasic cleavage site of HA was removed to a monobasic cleavage site, started losing body weight at 2 DPI (**Fig. 3a**) and succumbed to infection by 7 DPI (**Fig. 3b**). However, all mice infected with 10^2^, 10^3^, or 10^4^ PFU of LPhTXdNS1 maintained their initial body weight (**Fig. 3a**) and survived infection during the entire duration of the experiment (**Fig. 3b**). Consistently, viral replication of LPhTXdNS1 was significantly reduced *in vivo* when compared to LPhTX at 2 (**Fig. 3c**) and 4 (**Fig. 3d**) DPI in the nasal turbinate (NT) and lung homogenates. Unlike LPhTX, LPhTXdNS1 was not detected in brain tissues of infected mice (**Figs. 3c-d).**

**Figure 3.**
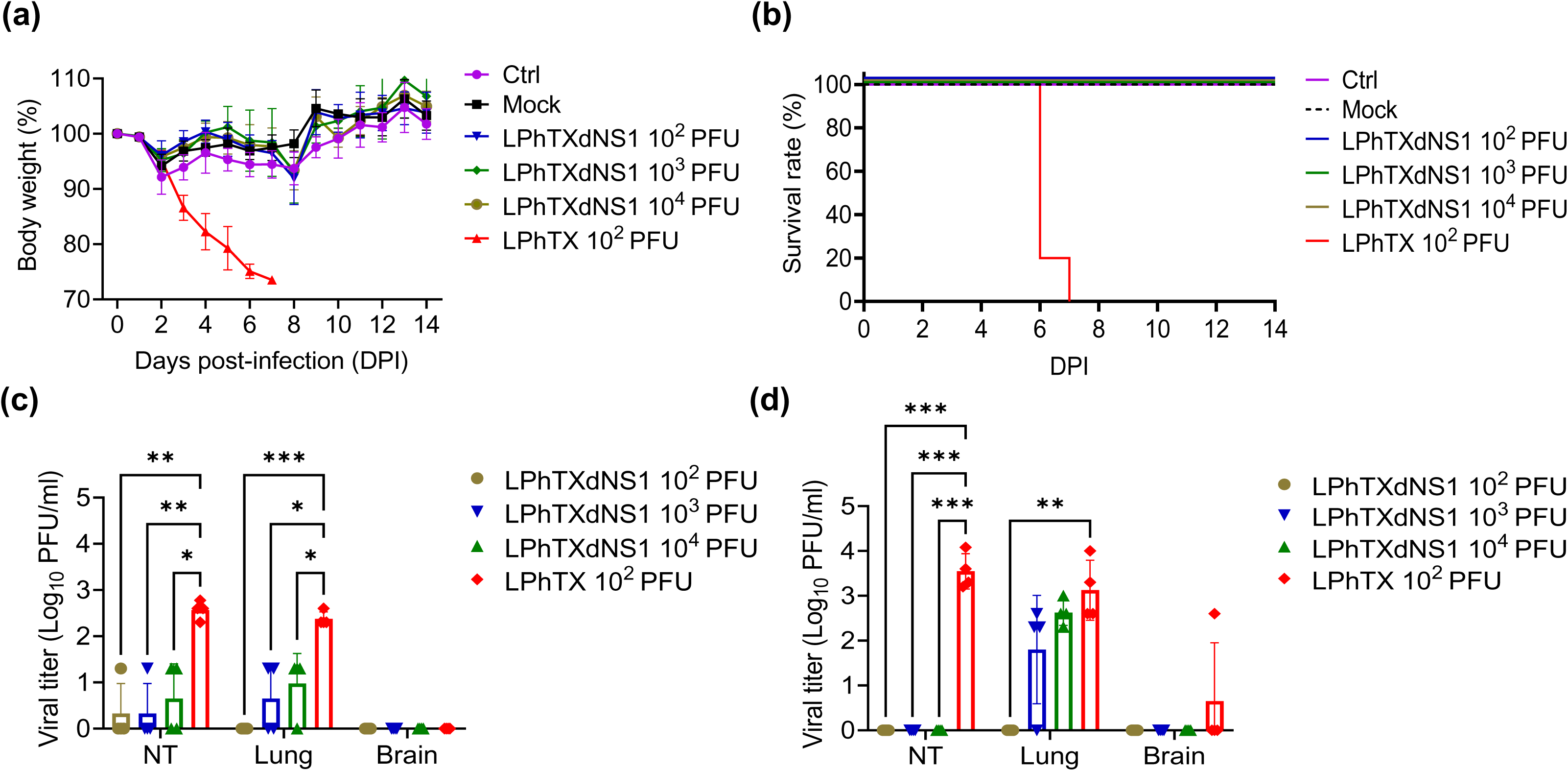
Pathogenicity of LPhTXdNS1. (**a**) Body weight loss of C57BL/6J mice infected with LPhTX (10^2^ PFU) or LPhTXdNS1 (10^2-4^ PFU), control (Ctrl), and mock-infected mice (inoculated with 1XPBS) (n=5). (**b**) Survival rates of C57BL/6J mice in panel a. (**c-d**) Viral replication of LPhTXdNS1 and LPhTX in the nasal turbinate (NT), lung and brain homogenates of mice at 2 (**c**) and 4 (**d**) DPI. Data are presented as mean ± SD. A two-way repeated measure ANOVA with Geisser-Greenhouse correction followed by Dunnett’s multiple comparisons test (* = *p* < 0.05, ** = *p* < 0.01, *** = *p* < 0.001).

To investigate immunogenicity, sera samples from LPhTXdNS1-infected mice were collected weekly for 4 weeks and assessed using HAI assay (**Table 3**). All mice infected with LPhTXdNS1 induced high levels of HAI titers against a highly pathogenic influenza A/Texas/37/2024 H5N1, designated hereafter as HPhTX, indicating its high immunogenicity even after a single dose of infection at lower (10^2^ PFU) dose (**Table 3**). Notably, we observed a direct correlation of HAI titers with the amount of LPhTXdNS1 used for vaccination.

**Table 3.** HAI titers against HPhTX in sera from mice immunized with LPhTXdNS1.

|  | HAI titer |  |  |  |  |  |  |  |  |  |  |  |  |  |  |
| --- | --- | --- | --- | --- | --- | --- | --- | --- | --- | --- | --- | --- | --- | --- | --- |
|  | LPhTXdNS1 10<sup>2</sup> PFU |  |  |  |  | LPhTXdNS1 10<sup>3</sup> PFU |  |  |  |  | LPhTXdNS1 10<sup>4</sup> PFU |  |  |  |  |
| **W1** | 20 | 10 | 20 | 10 | 10 | 40 | 40 | 20 | 20 | 20 | 80 | 40 | 80 | 20 | 20 |
| **W2** | 40 | 40 | 20 | 40 | 20 | 320 | 160 | 160 | 160 | 320 | 80 | 320 | 320 | 160 | 80 |
| **W3** | 160 | 80 | 160 | 320 | 80 | 320 | 640 | 320 | 160 | 320 | 320 | 640 | 320 | 640 | 320 |
| **W4** | 640 | 160 | 160 | 320 | 40 | 640 | 320 | 320 | 640 | 640 | 320 | 640 | 320 | 640 | 320 |

W: week

### Histopathology and immunohistochemistry (IHC) of LPhTXdNS1 infected mice

The lung and brain of infected mice were collected for pathological evaluation and quantification of viral antigen (**Fig. 4**). Pathological changes and high percentages of tissue viral antigen (NP) staining were detected in mice infected with 10^2^ PFU of LPhTX while mice infected with 10^2^ PFU of LPhTXdNS1 lack the presence of viral NP antigen, similar to mock-infected mice (**Figs. 4c-e**). As expected, mock-infected mice did not show any pathological lesions or viral antigen staining in the lungs and brain (**Figs. 4 a-e**). Mice infected with 10^2^ PFU of LPhTX showed the highest degree of pathological lesions (moderate to severe, multifocal to coalescing lymphoplasmacytic pneumonia) (**Figs. 4c and 5a**) and the highest area of lung staining for viral NP antigen (average percent pathology in the lung was approximately 22% (**Fig. 4c**) and average percentage of lung staining for viral antigen was approximately 19% (**Fig. 4d**)). Histological lesions in the lungs were characterized by multifocal to extensive areas of moderate alveolar damage frequently covered with fibrin and moderate infiltration of alveolar septa with lymphocytes, plasma cells and macrophages (**Figs. 4a and 5a**). Viral antigen staining was strongest in the bronchial epithelial cells that extended to surrounding alveolar epithelium and occasional nuclear staining of the cell in the alveolar septa (**Figs. 4b and 5b**). The average percent pathology in the lung was approximately ≤1%, and the average percentage of lung staining for viral NP antigen was approximately 5%, 0.7% and 0.1% for mice infected with 10^4^ PFU, 10^3^ PFU and 10^2^ PFU of LPhTXdNS1, respectively (**Figs. 4c-e**). No lesions in the brain were found in any of the LPhTXdNS1-infected mice, or LPhTX (**Figs. 4a and 5a**).

**Figure 4.**
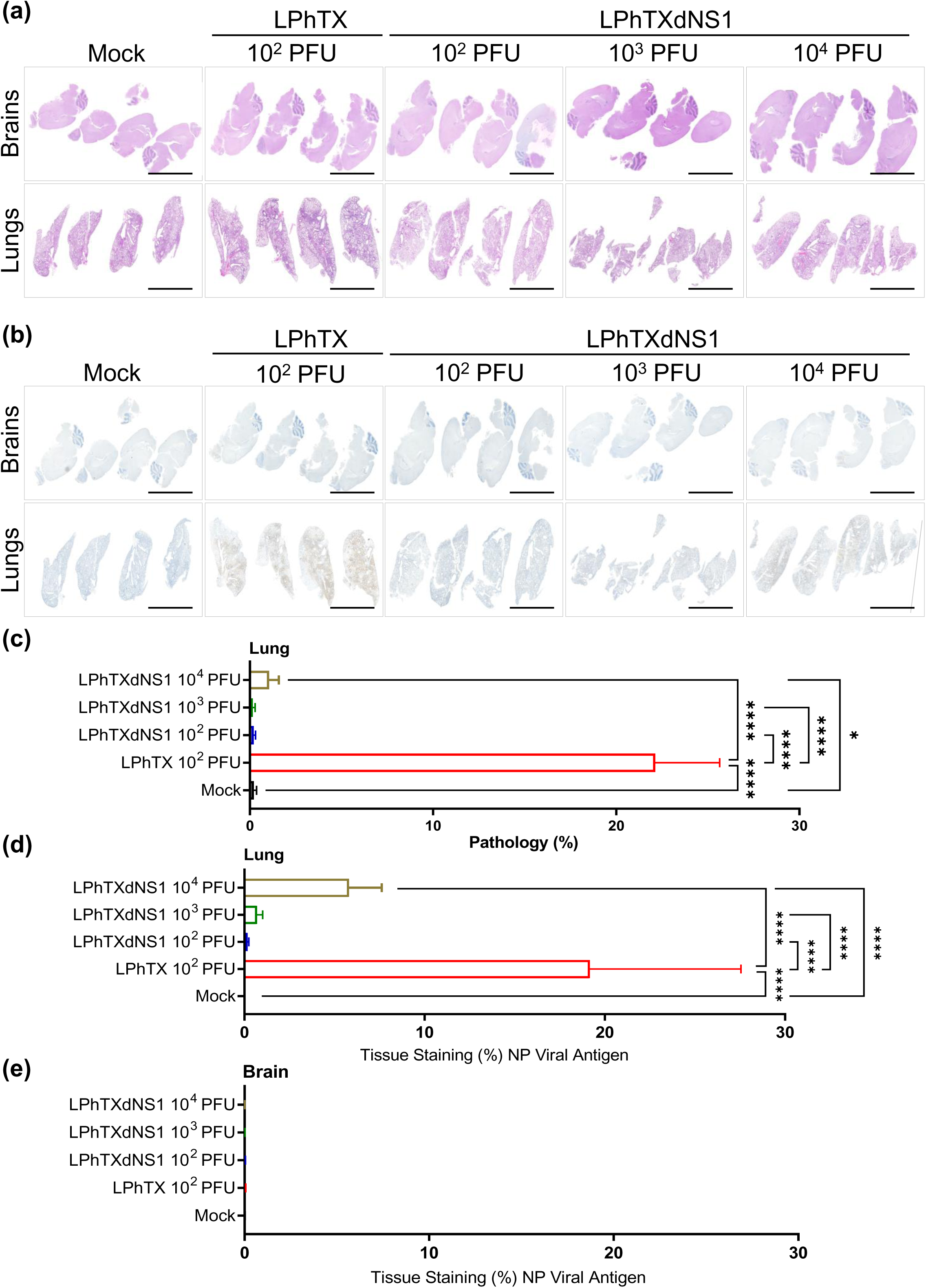
H&E and IHC staining of lung and brain tissues of mice infected with LPhTXdNS1. (**a**) H&E staining of brain (top) and lung (bottom) tissues of mock-infected and LPhTX- or LPhTXdNS1-infected mice. (**b**) Viral NP antigen IHC staining of brain (top) and lung (bottom) tissues of mock-infected and LPhTX- or LPhTXdNS1-infected mice. Scale bars represent 2 mm. (**c-e**) Quantitative assessment of histologic lesions (**c**) and tissue viral NP antigen positivity (%) in the lung (**d**) and brain (**e**) tissues from mock-infected and LPhTX- and LPhTXdNS1-infected mice. Data are presented as mean ± SD. Differences in quantitative assessment of histologic lesions and percentage of viral antigens were analyzed using beta regression. Post-hoc testing was conducted using Wald’s z-test with Tukey’s adjustment for multiple comparisons (* = *p* < 0.05; **** = *p* < 0.0001).

**Figure 5.**
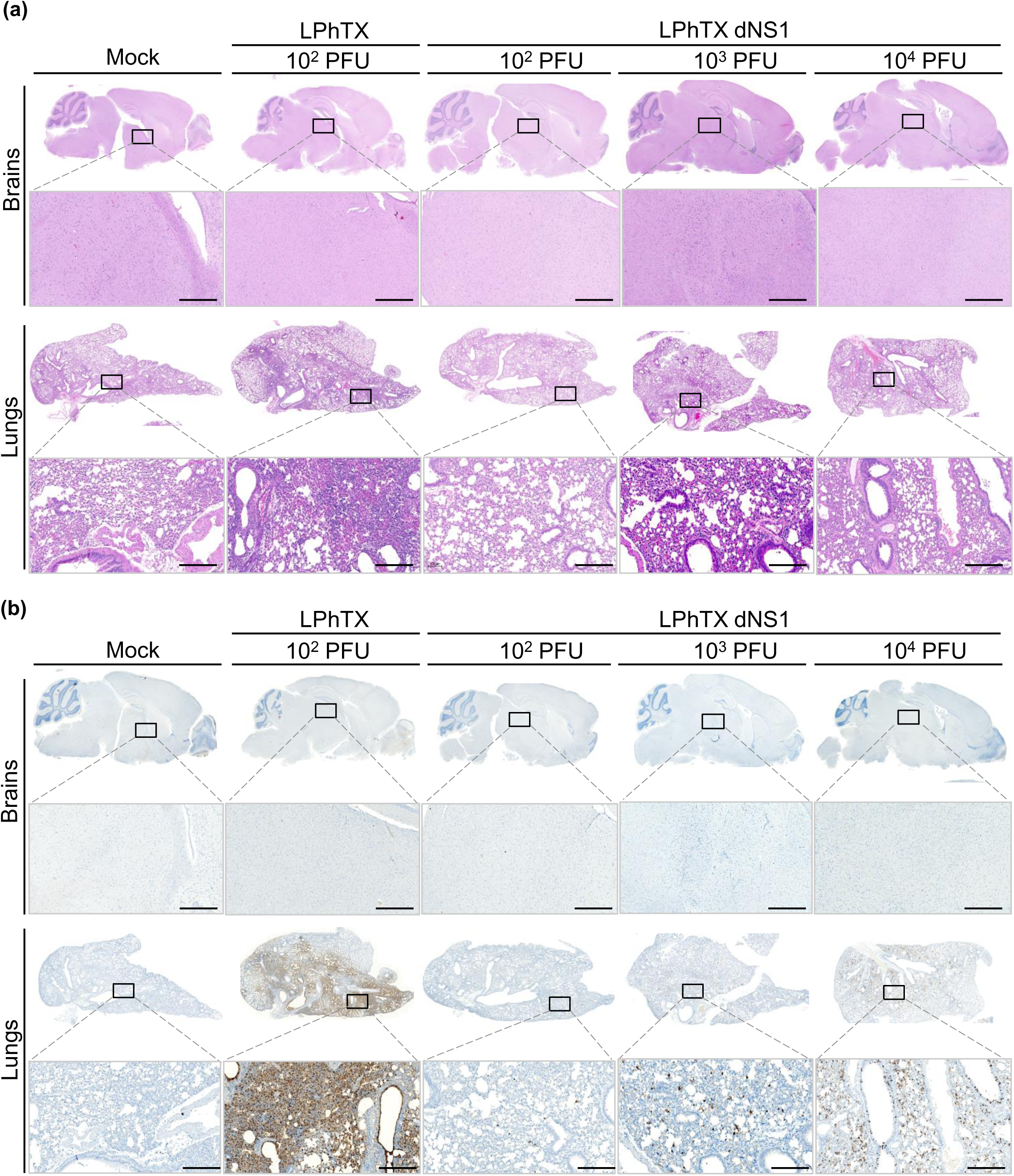
Histopathological findings in brain and lung sections of mice infected with LPhTXdNS1. H&E staining (**a**) and IHC detection of viral NP antigen (**b**) in the brain (top) and lung (bottom) tissues of mice mock-infected or infected with LPhTX and LPhTXdNS1. First row scale bars of each tissue represent 1000μm and second row scale bars of each tissue represent 100μm. The images displayed at the bottom are directly associated with the corresponding squares shown at the top images.

### LPhTXdNS1 vaccination protects against a lethal challenge with H5N1

Since infection with LPhTXdNS1 was not lethal in mice and was associated with seroconversion, we challenged LPhTXdNS1-infected mice with a lethal dose (10^3^ PFU) of a recombinant A/Texas/37/2024 H5N1 expressing nanoluciferase (Nluc), designated hereafter as HPhTX-Nluc (**Fig. 6**). As expected, mock-vaccinated mice challenge with HPhTX-Nluc lost body weight starting at 2 days post-challenge (**Fig. 6a**), and all of them succumbed to infection by days 6 and 7 after challenge (**Fig. 6b**). In contrast, all mice vaccinated with LPhTXdNS1 showed no reduction in body weight and all mice survived the lethal challenge with HPhTX-Nluc, except one mouse vaccinated with the lower dose of 10^2^ PFU of LPhTXdNS1 that succumbed to infection with HPhTX-Nluc at day 9 after challenge (**Figs. 6a** and **6b**). The presence of HPhTX-Nluc challenge virus in animals used to measure body weight changes and survival was detected using a non-invasive *in vivo* imaging system (IVIS). Strong Nluc signal in the lungs of mock-vaccinated mice was detected at days 4 and 6 post-challenge with HPhTX-Nluc (**Fig. 6c**), whereas no Nluc signal was detected in mice vaccinated with LPhTXdNS1, except in one mouse that was vaccinated with a lower dose of 10^2^ PFU of LPhTXdNS1 (**Fig. 6c**), that correspond to the mouse that did not survived challenge with HPhTX-Nluc (**Fig. 6b**). The serum samples for this mouse showed a reduced HAI titers when compared to other counterparts vaccinated with the same dose for up to 4 weeks post-infection (**Table 3**).

**Figure 6.**
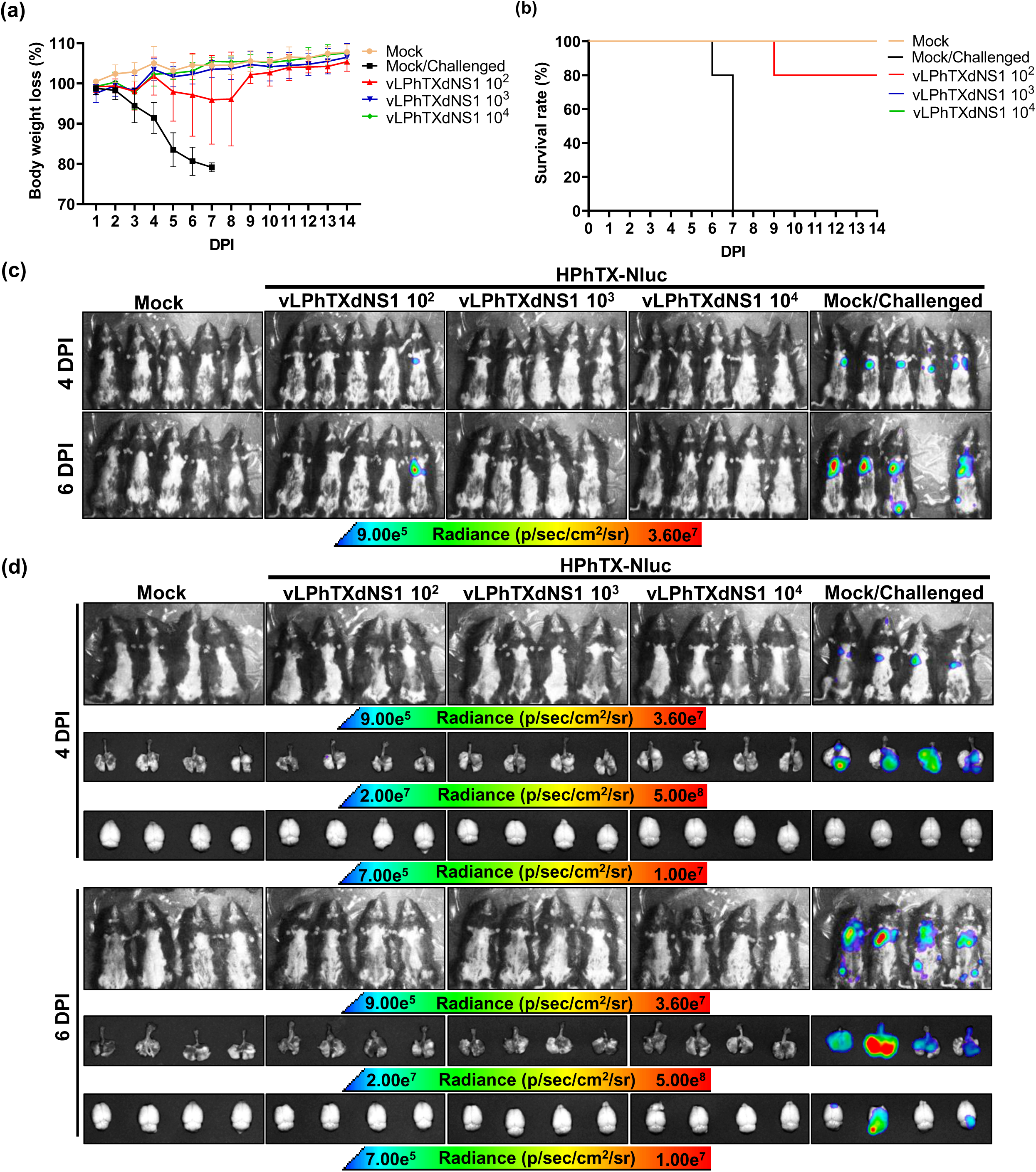
Protection efficacy of LPhTXdNS1. (**a**) Body weight loss (%) following vaccination with different doses of LPhTXdNS1 (vLPhTXdNS1) and challenge with 10^3^ PFU of HPhTX-Nluc. Mock-vaccinated animals challenge with 10^3^ PFU of HPhTX-Nluc, and mock-challenged mice, were included as control. (**b**) Survival rate following challenge with HPhTX-Nluc from animals in panel a. (**c**) Nluc expression as determined by IVIS from same group of animals used in panel a to assess morbidity and mortality at 4 and 6 DPI. (**d**) IVIS of necropsy groups of mice and their corresponding lung and brain tissues at 4 and 6 DPI.

Consistently, in the necropsy mice groups, strong Nluc signal was detected in the lungs of mock-vaccinated mice at days 4 and 6 post-challenge with HPhTX-Nluc (**Fig. 6d**), whereas no Nluc signal was detected in LPhTXdNS1-vaccinated mice (**Fig. 6d**). All vaccinated mice groups, including those with the lower dose (10^2^ PFU) of LPhTXdNS1 did not show Nluc signal in the lungs or brains (**Fig. 6d**). Notably, we were able to detect Nluc signal in the brain of mock-vaccinated mice after challenge with HPhTX-Nluc at day 6 post-infection, although only in 2 of the challenged animals (**Fig. 6d**). The protection efficacy of LPhTXdNS1 was also confirmed by assessing viral titers of HPhTX-Nluc in the NT, lungs, and brains of vaccinated animals by viral titration, and Nluc expression (**Fig. 7**). As expected, we detected high viral titers of HPhTX-Nluc in mock-vaccinated mice as determined by Nluc expression (**Figs. 7a and 7c**) and viral titration (**Figs. 7b and 7d**) at 4 (**Figs. 7a and 7b**) and 6 (**Figs. 7c and 7d**) days post-challenge with HPhTX-Nluc. These *in vivo* results indicate the high protection efficacy of LPhTXdNS1 against HPhTX.

**Figure 7.**
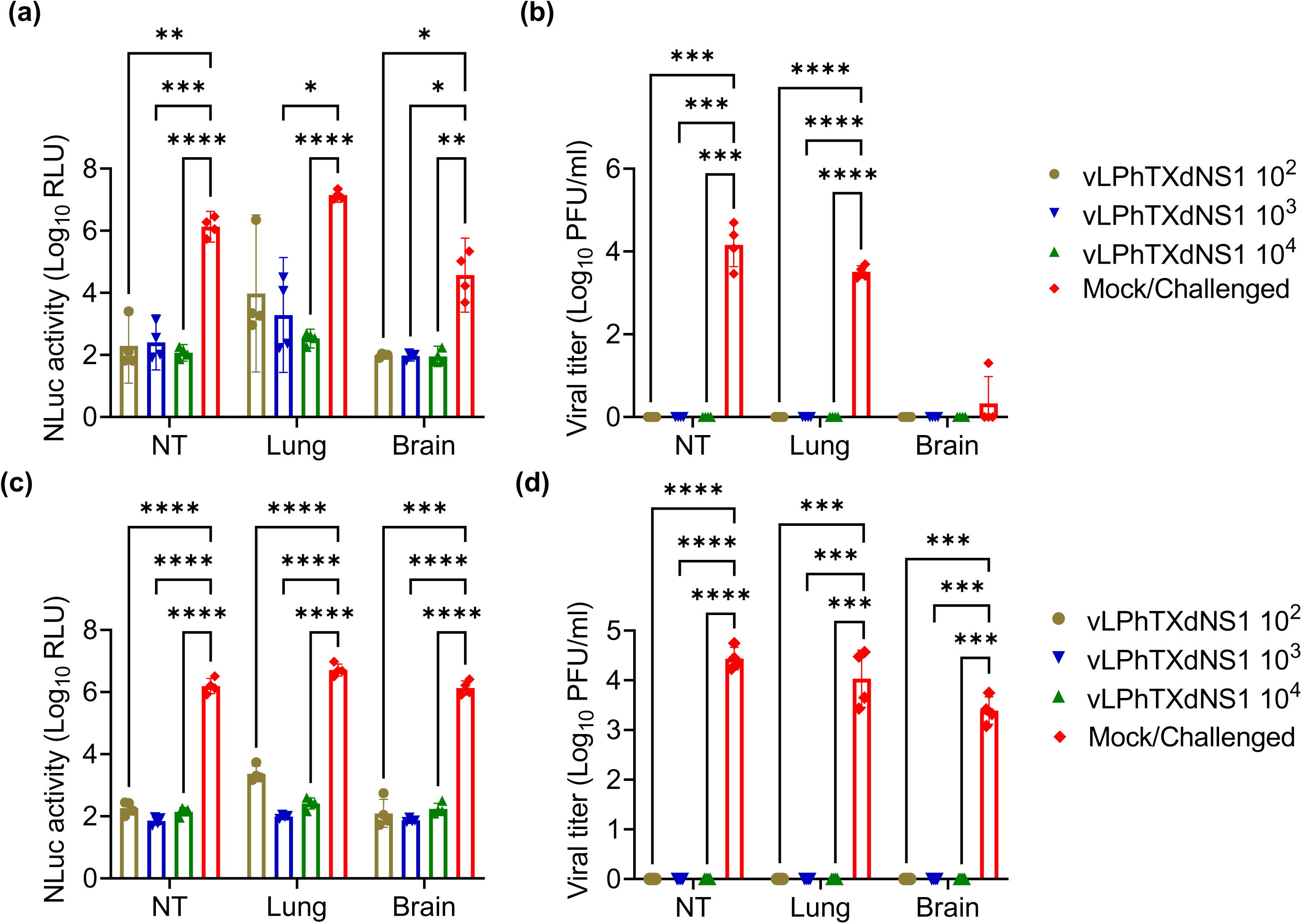
HPhTX-Nluc viral titers in the NT, lungs, and brains of mice vaccinated with LPhTXdNS1. (**a & b**) Viral replication of HPhTX-Nluc was determined by measuring Nluc activity (**a**) and viral titers (**b**) at day 4 post-challenge. (**c & d**) Viral replication of HPhTX-Nluc was determined by Nluc activity (**c**) and viral titers (**d**) at day 6 post-challenge. Data are presented as mean ± SD. A two-way repeated measure ANOVA with Geisser-Greenhouse correction followed by Dunnett’s multiple comparisons test (* = *p* < 0.05, ** = *p* < 0.01, *** = *p* < 0.001, **** = *p* < 0.0001).

### Histopathology and IHC of mice vaccinated with LPhTXdNS1 following challenge with HPhTX-Nluc

The protective efficacy of LPhTXdNS1 was further confirmed by evaluating histopathological lesions and the presence of viral NP antigen in the lung and brain tissues of vaccinated mice after challenge with HPhTX-Nluc (**Fig. 8**). **Figures 8a and 8b** represent the H&E and IHC stained brain and lung tissues of all mice included in the study, respectively. **Figures 9a and 9b** show the H&E and IHC stained brain and lung of representative mice at higher magnifications. HPhTX-Nluc- challenged, mock vaccinated, mice showed the most severe pathological lesions, that were similar to those seen in animals infected with 10^2^ PFU of LPhTX, consisting of moderate to severe, multifocal to coalescing lymphoplasmacytic pneumonia (**Figs. 8a and 9a**). The average percent pathology in the lung was approximately 37% (**Fig. 8c**) and average percentage of lung staining for viral antigen was 7% (**Fig. 8d**). One mouse in this group showed multifocal, mild perivascular cuffing of lymphocytes in the mid brain (**Fig. 9a**, mild multifocal lymphoplasmacytic encephalitis). LPhTXdNS1-vaccinated mice challenged with HPhTX-Nluc showed mild to minimal pathological lesions. The lesions and IHC pattern were comparable to those in mock-infected mice (**Fig. 9a**). The average percent pathology in the lung was ≤1% and average percentage of lung staining for viral antigen was ≤1% for all vaccinated mice with the different doses of LPhTXdNS1, respectively. There were multifocal areas of viral NP staining in the mid brain and brain stem regions (approximately 3%) (**Figs. 8e and 9b**) and scattered viral NP staining in bronchial epithelial cells and in the alveolar septa of infected mock-vaccinated mice (**Figs. 9a and 9b**). The vaccinated groups showed rare viral NP staining in the bronchial epithelial cells and alveolar septa. No viral NP antigen was detected in the brain of LPhTXdNS1-vaccinated, HPhTX-Nluc-challenged groups (**Fig. 9b**).

**Figure 8.**
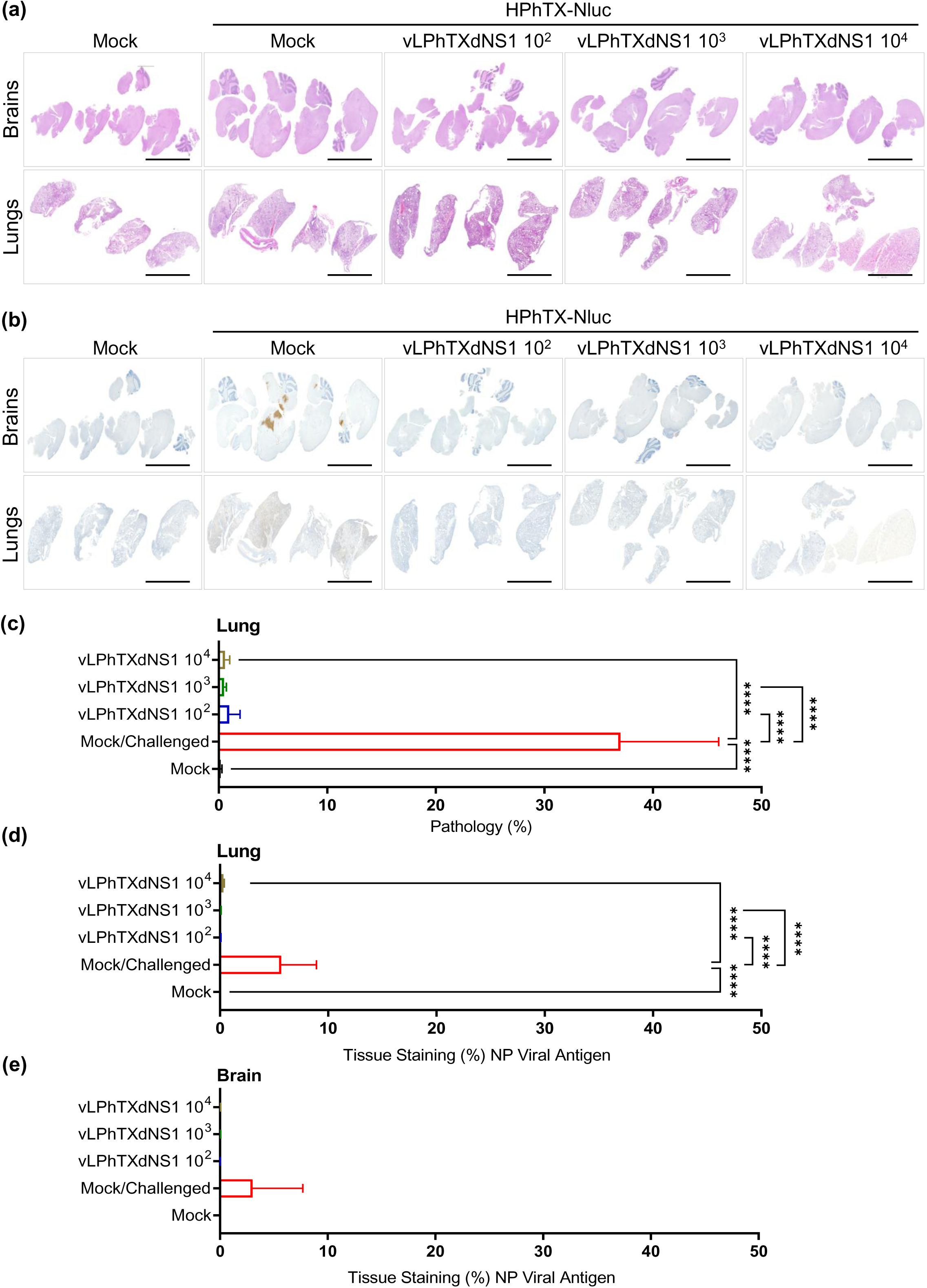
Histologic lesions and IHC staining for viral NP antigen after challenge with HPhTX-Nluc. (**a-b**) H&E tissue sections (**a**) and IHC (**b**) of brain (top) and lung (bottom) tissues from mock-vaccinated, mock-challenged mice; mock-vaccinated, challenged mice; and vaccinated mice (10^2-4^ PFU vLPhTXdNS1), challenge mice. All challenged groups were infected with a lethal infection dose of 10^3^ PFU of the HPhTX-Nluc. Scale bars represent 2 mm. (**c-e**) Quantitative assessment of histologic lesions (**c**) and tissue viral NP antigen positivity (%) in the lung (**d**) and brain (**e**) tissues from mice in panel a. Data are presented as mean ± SD. Differences in quantitative assessment of histologic lesions and percentage of viral antigens were analyzed using beta regression. Post-hoc testing was conducted using Wald’s z-test with Tukey’s adjustment for multiple comparisons (**** = *p* < 0.0001).

**Figure 9.**
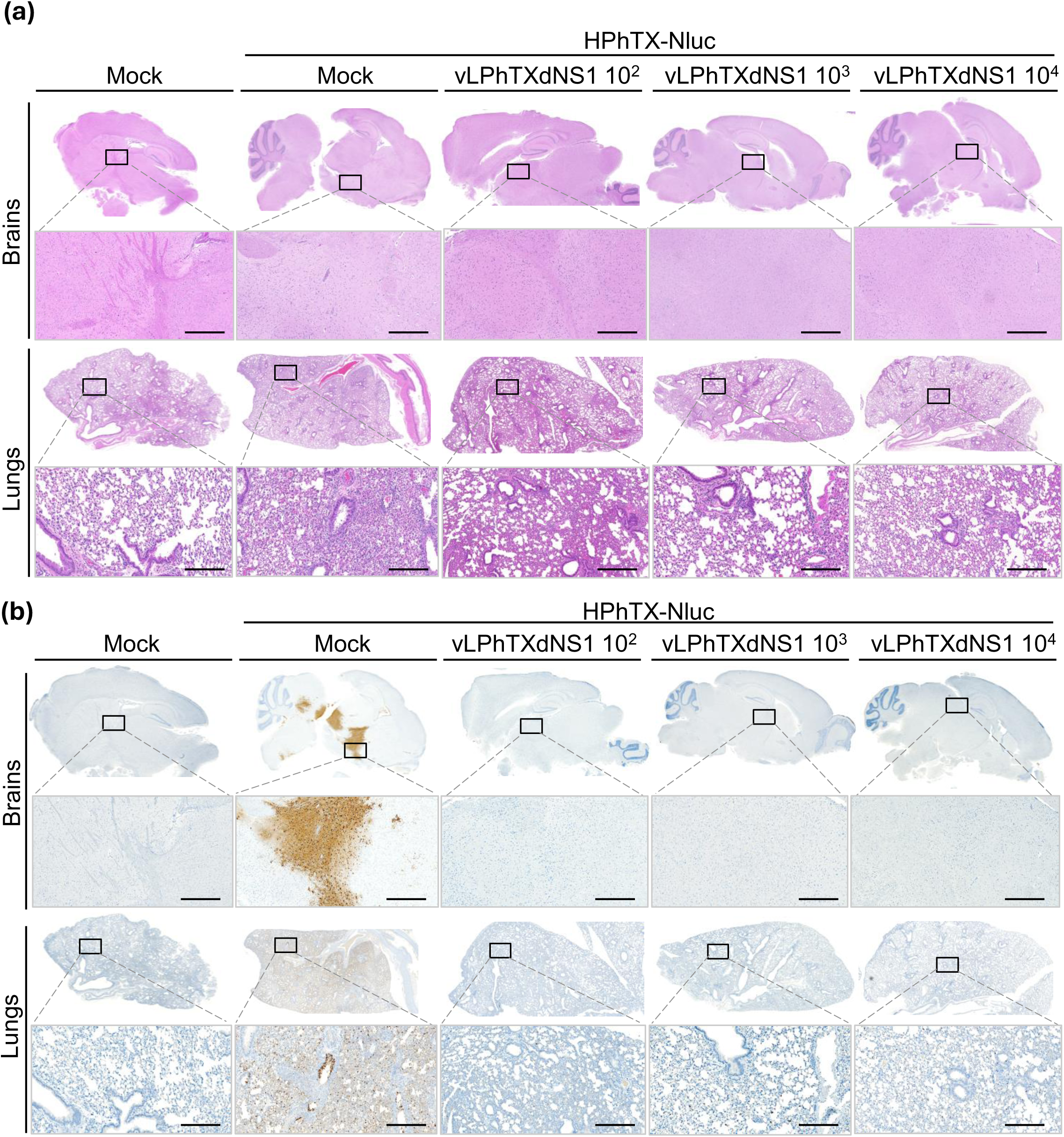
Histopathological findings in the brain and lung sections of mice challenge with HPhTX-Nluc. (**a**) H&E staining and (**b**) IHC detection of viral antigen (NP) in in the brain (top) and lung (bottom) tissues. First row scale bars of each tissue represent 1000μm and second row scale bars of each tissue represents 100μm. The images displayed at the bottom are directly associated with the corresponding squares shown at the top images.

## Discussion

Influenza is one of the most significant respiratory viral pathogens, responsible for about 2% of all annual respiratory deaths (42). It predominantly affects high-risk groups, including children, the elderly, pregnant women, and immunocompromised individuals (43). Vaccination is the most effective methods for preventing influenza disease, especially in high-risk groups (43). Along with inactivated influenza vaccines, LAIVs play a crucial role in preventing seasonal IAV infections. LAIVs offer the advantages of needle-free administration, the ability to stimulate mucosal immune responses, and are approved for use in healthy individuals aged 2 to 49 years (8, 44).

The currently approved LAIV is based on a genetically modified, *ts*, *ca*, *att* influenza strain derived from A/Ann Arbor/6/1960 H2N2 with restricted ability to replicate in the upper (33°C) rather than lower (37°C) respiratory tract, preventing disease onset. LAIV mimics a natural infection, offering the advantage of stimulating mucosal, cellular, and cross-protective immune responses. Despite these advantages, the effectiveness of the LAIV in many studies is comparable, or even lower, than the inactivated influenza vaccine. Besides, the possible reversion of the changes that are responsible for the ts, ca, att of the current form of LAIV due to the evolutionary process may enhance virus replication (45). Unexpected replication of the vaccine virus and reversion to pathogenic strains in immunocompromised patients has been documented for other attenuated vaccine strains (46–48). Therefore, certain key populations are excluded from LAIV recommendations.

The current outbreak of HPAIV H5N1 clade 2.3.4.4.b, genotype B3.13, among dairy cattle was first identified in the US in March 2024 (1). In parallel, the genotype D1.1, a reassortant of genotype A3, could quickly spread to all North American flyways and became the predominant genotype in migratory birds, and was recently also documented in dairy cattle in Nevada with adaptive D701N and E627K markers of mammalian host adaptation (3). Human cases of HPAIV H5N1 among dairy farm workers started to be reported in April 2024 in Texas. Cumulatively, 70 confirmed HPAIV H5N1 cases (clade 2.3.4.4b, genotypes B3.13 and D1.1) have been documented including one fatality (49). While the current public health risk remains low, these findings have increased the potential for exposure to the virus among individuals working with infected poultry and cattle, such as farm and dairy workers and veterinarians. As a result, the development of an effective HPAIV H5N1 2.3.4.4b vaccine has become essential to protect these high-risk groups.

In this study, we developed an efficient and safe live-attenuated dNS1-based 2.3.4.4b H5N1 candidate vaccine with high immunogenicity and efficacy in mice. Unlike the currently in-use LAIVs, which incorporate changes to make the vaccine strain ts (50), the complete deletion of the NS1 protein ensures that the virus is unlikely to restore pathogenicity after successive passaging (51). Several studies have explored the use of truncated or deleted NS1 as a safe vaccination platform against various IAVs, including those affecting avian (52), swine (53), canine (54), equine (55), and humans (22, 56), as well as H5N1 (57), demonstrating its high safety profile. Moreover, a recent study suggested a mechanism to minimize the risk of the current LAIV by applying a targeted approach to mutate the NS1 protein of the LAIV backbone making it unable to bind the host factor CPSF30 (58), allowing the processing of host pre-mRNAs, including those for IFN (21). Consistently, in this work, infections with the highly pathogenic (HPhTX) and low pathogenic (LPhTX) strains with intact NS1 were associated with systemic infection, including neurovirulence in the case of HPhTX (25), while infection with LPhTXdNS1 did not result in systemic infection and/or neurotropism. This emphasizes that LPhTXdNS1 could be a safer option as a LAIV.

One potential concern with using LPhTXdNS1 is the risk of reassortment with other IAV that may naturally infect the vaccinated individual, potentially leading to the emergence of new virus reassortants. However, reassortment typically occurs when two IAVs with efficient replication co-infect the same host cell (59). The deletion of NS1 in H5N1 viruses typically leads to a virus that induces a stronger innate immune response, which can restrict its ability to replicate efficiently, or block the infection of other influenza strains that are sensitive to IFN. This reduces the severity of the infection and make the virus less capable of spreading in the host (60). It is also important to note that LPhTXdNS1 exhibits reduced replication efficiency in standard immunocompetent mammalian cells, and it has limited replication outside IFN-deficient Vero cells, which are recommended for growing IFN-sensitive viruses, including IAV lacking NS1. These characteristics reduce the likelihood of reassortment of LPhTXdNS1 with other IAVs.

Taken together, we have developed LPhTXdNS1 and demonstrated its potential as a safe, immunogenic, and effective vaccine against the recently emerged 2.3.4.4b H5N1. This work serves as a proof-of-concept study for the development of a universal avian influenza vaccine targeting currently emerging strains with potential zoonotic risks. We propose that further studies are needed to evaluate the impact of a multidose LAIV approach and to investigate the vaccine’s efficacy in animals, including those currently susceptible to HPAIV infections where the LPhTXdNS1 vaccine could be used (poultry and farm animals). Our approach offers the potential to expand the applications of the current LAIV, particularly in targeting high-risk populations where the burden of IAV remains a significant concern.

## Acknowledgments

We thank the Histology unit team staff, Dr. Renee Escalona and Mr. Colin Chuba, for their assistance in tissue staining/immunostaining experiments, and the Cell Biology Core Lab at Texas Biomedical Research Institute for assistance with the multiplex cytokine assay. We also acknowledge the BEI Resources for providing the PB1 (Clone F5-46) and PA (Clone 1F6) monoclonal antibodies.

## Authors Contributions

Conceptualization: A.M. and L.M-S.; Methodology: A.M., C.Y., R.S.B., V.S., R.M., R.N.P., R.A.E., M.B., E.M.C., N.J., A.C., and A.N.; Data collection and interpretation: A.M., V.S., R.M., T.J.C.A., C.Y., A.G-S., and L.M-S.; Funding acquisition and resources: A.M., T.J.C.A., A.G-S., and L.M-S.; Writing-original draft preparation: A.M. and L.M-S.; Writing-review and editing: all authors have read and agreed to the published version of the manuscript.

## Data availability

All data generated or analyzed during this study are included in this published article

